# Evaluation of SNP-based genotyping to monitor tuberculosis control in a high MDR-TB setting

**DOI:** 10.1101/044370

**Authors:** N Tukvadze, I Bergval, N Bablishvili, N Bzekalava, ARJ Schuitema, J de Beer, R de Zwaan, S Alba, D van Soolingen, R Aspindzelashvili, RM Anthony, S Sengstake

## Abstract

*Mycobacterium tuberculosis* (*Mtb*) lineage identification and typing of clinical isolates in general is performed only retrospectively. The results are rarely linked to drug susceptibility testing (DST) or patient data. Consequently, the association between Mtb lineage, (multi)drug resistance and treatment history is not fully explored at the local level. Here we evaluated a new SNP based typing assay. We furthermore assessed the added value of genotyping of Mtb isolates for epidemiological purposes and guidance of tuberculosis (TB) control. *Mtb* lineage, DST profile and treatment history were determined for 399 samples at the National TB Reference Laboratory (NRL) in Tbilisi, Georgia by local staff. Data was shared electronically and analysis was performed remotely. Out of 399 isolates, 74 (74/399, 18.5%) were at least multidrug resistant (MDR)-TB, of which 63 (63/74, 85.1%) were members of three different *Mtb* Beijing lineages. Previous treatment was reported in 38/74 (51.4%) MDR(+) patients. The availability of this data allows associations with lineages. Notably, multidrug resistant TB was more strongly associated with the Beijing lineage than treatment history. Of all MDR-TB Beijing strains 56.7% (42/74) were members of a genetic cluster. This is most easily explained by (ongoing) MDR-TB transmission rather than drug resistance amplification. This knowledge is useful when designing intervention strategies for MDR-TB. Our study provides an example that on-site integrated *Mtb* genotyping is realistic and could support TB control activities.

## INTRODUCTION

The WHO has approved a post-2015 Global End Tuberculosis Strategy for tuberculosis (TB) prevention, care and control (1). Countries need to respond by adapting and enhancing their TB control activities (1, 2). Justifying investment in effective TB control strategies in a country can be achieved in part by defining and monitoring the (MDR) TB epidemic to identify appropriate interventions.

Molecular tools can positively impact on earlier detection of *Mtb* and identification of drug resistance (3, 4). Genotyping of *Mtb* isolates has revealed associations between drug resistance and *Mtb* lineage (5–8), identified routes of transmission (9, 10) and described the dynamics of epidemic clones (3, 11–14). Further developments in multiplex assays as well as the expanded use of next generation sequencing assays will increasingly allow *Mtb* strains to be simultaneously screened for resistance associated mutations and the bacterial lineage they represent.

A robust link has been found between previous treatment for TB and multidrug resistance (15), and is identified as a risk factor for MDR-TB by the WHO (16) but other factors are also important, for example the bacterial lineage. This is especially true when transmission of resistant strains is more common than the acquisition of resistance during treatment. Members of the East Asia lineage (*Mtb* lineage 2) (17, 18) have repeatedly been associated with multidrug resistance in high burden MDR-TB countries (11, 19) but less so in low burden (MDR)-TB countries (20–22). The relative importance and interdependence of these factors for infection control has received comparatively little attention.

Georgia is a high burden MDR-TB country with 17.7% MDR-TB and 3.3% extensively drug resistant (XDR)-TB reported in 2013 (23). Georgia's geographical setting between Eastern Europe, Russia and East-Asia is reflected in the genetic diversity of circulating *Mtb* strains (5, 24). Prior to this study there was no local capacity in Georgia to routinely and prospectively identify, document or monitor the genotypes of isolated *Mtb* strains. Previous studies have shown that in Georgia the Beijing lineage is associated with multidrug resistance (5, 24, 25).

Here, we evaluated the performance of a SNP-based molecular assay for *Mtb* genotyping and especially its practicality and value when linked to patient data and phenotypic DST at the NRL in Tbilisi, Georgia. The combined data provide an insight into the dynamics of infection and the feasibility of genotyping as a routine component of a national TB reference laboratory. Our data suggest that monitoring and interrupting the spread of Beijing genotype MDR-TB clones is of the utmost importance. Strengthening TB infection control by ongoingmonitoring of the circulating genotypes can provide data to support continued investment in these activities.

## MATERIAL and METHODS

### Patient material

Between August 2012 and April 2013, 30.5% of all well grown diagnostic cultures from individual pulmonary TB patients (a total of 399 samples) were randomly selected each month (approximately 40 per month) for analysis at the National TB Reference Laboratory in Tbilisi, Georgia. Patient samples included those from patients administered directly at the NCTLD in Tbilisi and also from the nine country-wide microscopic centers.

Informed consent was not required as the patient information used was anonymized before linking to the results of the analysis of the bacterial cultures and could not be linked back to individual patients.

#### Patient data

Anonymized patient data (age, patient treatment status, patient outcome, DST, molecular resistance testing) were extracted from the patient database at the NRL and communicated to the KIT for further analysis.

#### DNA extraction

DNA was extracted on site at the NRL in Tbilisi by thermolysis and sonication according to the Genotype MTBDR*plus* protocol (Hain, Nehren, Germany).

### MLPA assay

A total of 399 DNA samples were analyzed by Multiplex Ligation-dependent Probe Amplification (MLPA) using xTAG technology on a MAGPIX^TM^ device (Luminex BV, Austin, Texas, USA) as previously described (26, 27) in 10 runs in Tbilisi, by local laboratory staff after one week of onsite training. In each run eight or more of the cultures were from a sputum smear negative case. Data from each run was emailed in the form of a csv file for remote analysis.

MLPA profiles were assigned on the basis of the calculated values of previously published markers (24) and newly added validated MLPA oligos targeting the eisG-10A and eisG-14T mutation (eisG10-LPO 5’-CGTGGCCGCGGCATATGCCACAA-3’ and eisG10-RPO 5’-TCGGATTCTGTGACTGTGACCCTGTGTAGCCCGACCGAGGACGACTGGCC-3’ eisG14-LPO 5’-TCAGGGTCACAGTCACAGAATCCGACTGTA-3 and eisG14-RPO 5’-GCATATGCCGCGGCCACGTGCACGTGAATATTACGACGACAGTGTCTGG-3’). Intermediate marker values for drug resistance targeting probes were interpreted as heteroresistance of the respective allele (28). Lineage identification by MLPA was performed by targeting lineage specific markers described previously (26).

### MLPA data analysis

Briefly all data obtained from the csv files of the individual MAGPIX runs received in Amsterdam were combined and analyzed in dedicated excel sheets as previously described (24). Intra-normalization was performed on the raw Median Fluorescence Intensity (MFI) signals followed by the application of marker-specific correction factors (24).The default range for intermediate values was defined between a corrected MFIof 330-590. After this analysis the average number of intermediate values per strain was just below 1 (0.80). Using the sigmoid curves generated from the data set to adjust the corrected MFI range the number of intermediate values per strain was further reduced to 0.35 ((24), Figure 2B). This data was linked to DST and patient information collected in Georgia. Any intermediate calls for drug resistance markers were regarded as resistant by MLPA and assumed to represent mixed genotypes.

### Phenotypic and molecular drug resistance detection

Phenotypic DST and GenotypeMTBDR*plus* (hereafter, MTBDR*plus*) were routinely performed by the staff at the NRL (3) and results were anonymized, documented in electronic data files and sent to the KIT.

### Sequencing

PCR amplification and sequencing of the *embB, gyrA* genes in selected isolates was performed to verify the MLPA results with the following primers: gyrA and embB (26) Sequencing of PCR products was performed by Macrogen Inc. (Amsterdam, The Netherlands).

### MIRU-VNTR typing

An optimized version (29) of the standard VNTR typing using 24 loci (30) was performed at the RIVM at the RIVM, Bilthoven, the Netherlands. Identification of MLVA 15-9 codes was carried out by using the MIRU-VNTRplus database (31). A cluster was defined as a minimum of two isolates with identical MIRU-VNTR patterns.

### Statistical analysis

Analysis of sensitivity, specificity, PPV and NPV of the MLPA in comparison to DST and the MTBDR*plus* assay was performed using GraphPadPrism version 5.03. The kappa coefficient was calculated using GraphPadPrismQuickCalcs (http://www.graphpad.com/quickcalcs/). Univariate and multivariate regression analysis was performed using STATA statistical software, Breda, The Netherlands.

## RESULTS

After initial automated MLPA data analysis (24), 43 of the 399 strains were not automatically assigned to a lineage and required expert review. After this process 388 (97.2%) of the samples were assigned to a single lineage; 32 after expert review. Of the remaining 11 strains, five remained uninterpretable and six were identified as having a mixed profile consistent with the presence of two lineages.

An overview of interpretable results obtained by each method (MLPA, DST, GenotypeMTBDR*plus*) is summarized in FIGURE 1. DST identified an MDR-TB phenotype in 74/399 (18.5%) patient samples (TABLE 1). Of these, eight (10.8%) strains were identified as XDR-TB.

**Figure 1.**
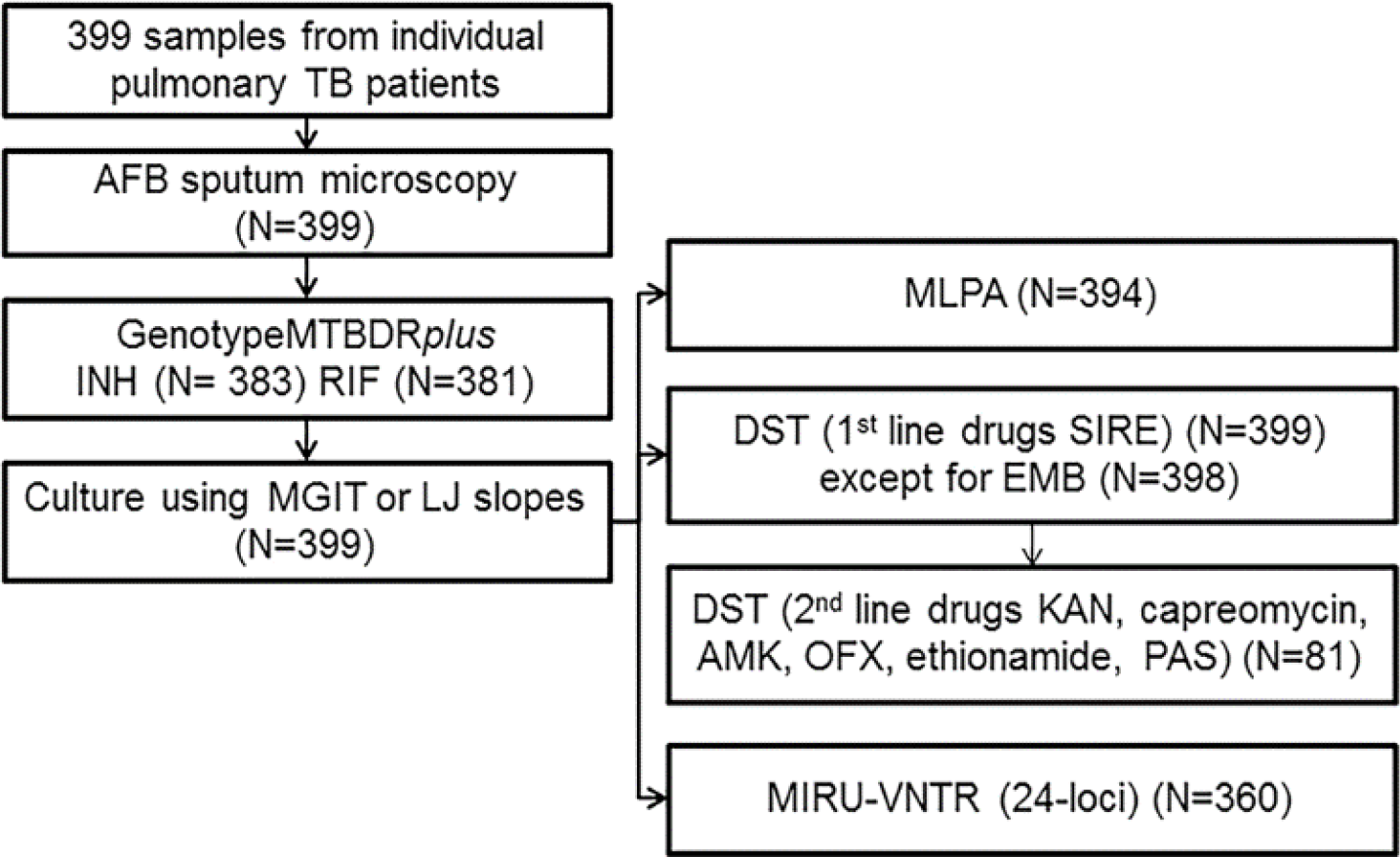
Overview of interpretable results obtained by each method. STR-streptomycin, INH-isoniazid, RIF-rifampicin, EMB-ethambutol, KAN-kanamycin, CAM-capreomycin, AMK-amikacin, OFX-ofloxacin, ETH-ethionamide, PAS-para aminosalicylic acid. DST for the first line drugs STR, INH, RIF and EMB and the second line drugs ETH, PAS, KAN, CAM and OFX was performed at the NRL as described elsewhere (3, 44). Molecular resistance testing and confirmation of Mycobacterium tuberculosis complex was performed directly on sputum samples and/or on cultures using the Genotype MTBDR*plus* assay (44, 45) at the NRL. 24-locus MIRU-VNTR typing (29) was performed either at the RIVM or by Genoscreen(Lille, France)). DST results for first line drugs resistance were obtained from all 399 isolates. DST for second line drug resistance was performed on 82 isolates, valid results were obtained for 81 isolates. Interpretable MLPA profiles were obtained from 394 (99.0%) strains. Using the MTBDR*plus* assay interpretable results for isoniazid and rifampicin resistance were obtained for 383/399 (96.2%) and 381/399 (95.7%) strains, respectively.

**TABLE 1.**
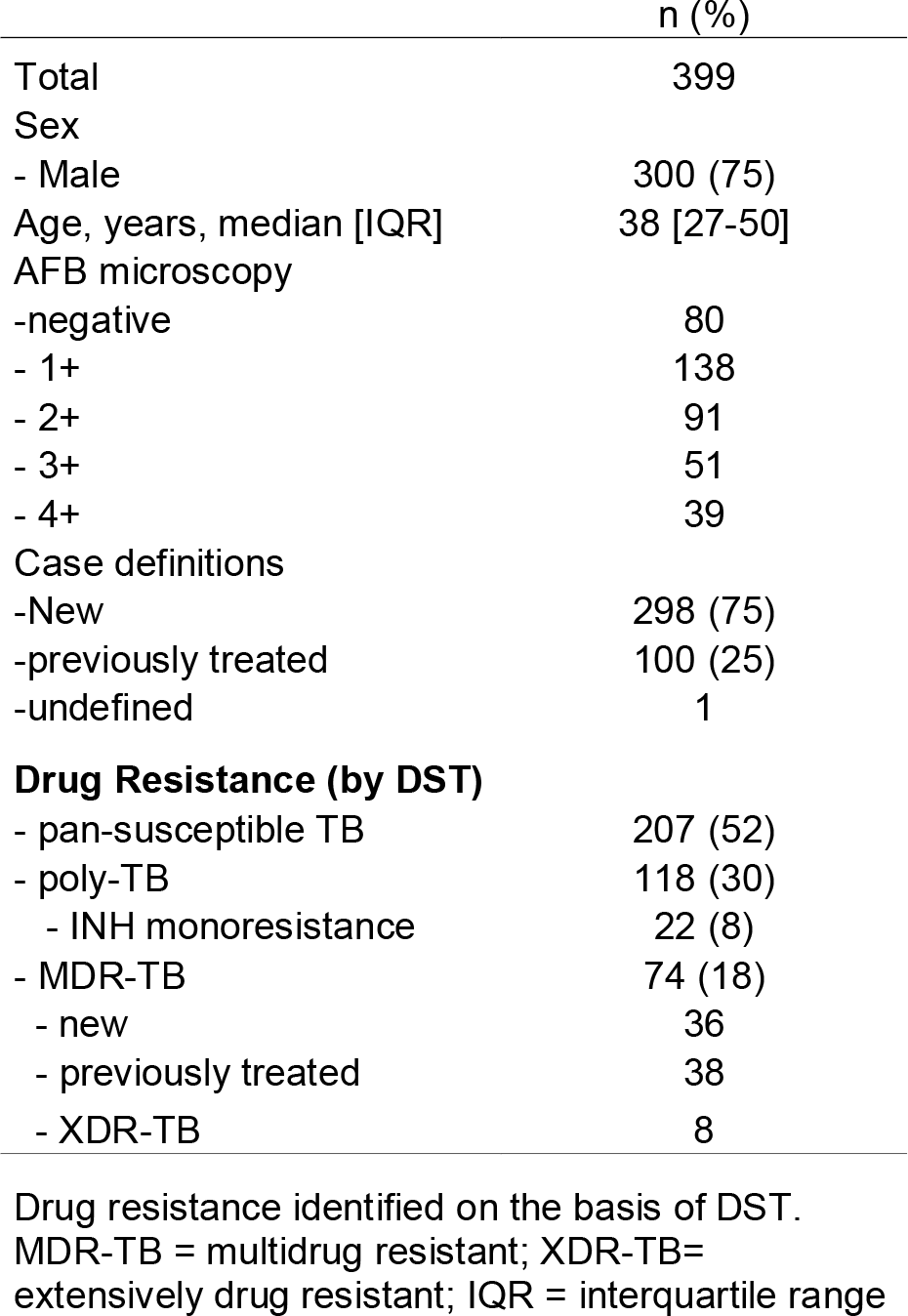
Baseline characteristics of all patients enrolled and drug resistance identified

|  | n (%) |
| --- | --- |
| Total | 399 |
| Sex |  |
| - Male | 300 (75) |
| Age, years, median [IQR] | 38 [27-50] |
| AFB microscopy |  |
| -negative | 80 |
| - 1+ | 138 |
| - 2+ | 91 |
| - 3+ | 51 |
| - 4+ | 39 |
| Case definitions |  |
| -New | 298 (75) |
| -previously treated | 100 (25) |
| -undefined | 1 |
| **Drug Resistance (by DST)** |  |
| - pan-susceptible TB | 207 (52) |
| - poly-TB | 118 (30) |
| - INH monoresistance | 22 (8) |
| - MDR-TB | 74 (18) |
| - new | 36 |
| - previously treated | 38 |
| - XDR-TB | 8 |

Drug resistance identified on the basis of DST. MDR-TB = multidrug resistant; XDR-TB= extensively drug resistant; IQR = interquartile range

DST and MTBDR*plus* confirmed 313 of 344 resistance associated mutations identified by MLPA, for 12 of the 344 MLPA detected mutations there was no valid data available by either DST or MTBDR*plus*. An intermediate marker value by MLPA was obtained for 28 (8.9%) of the 313 resistance MLPA calls. The 12 (3.8%) MLPA resistance calls not supported by DST or MTBDR*plus* all had intermediate values. Six of these 12 intermediate resistance calls were for RIF resistance associated mutations for which the MTBDR*plus* assay identified the wild type sequence only (data not shown). Tables showing sensitivity and specificity values for drug resistance detection by MLPA compared to DST (TABLE A1) and MTBDR*plus* assay (TABLE A2) are provided as supplementary information.

RIF resistance was conferred by the rpoB-531 mutation in more than half of all MDR-TB strains based on MTBDR*plus* (41/66; 62.1%) and MLPA (51/55; 96.4%) results (TABLE A3). MTBDR*plus* identified RIF resistance based on the loss of an *rpoB* wildtype probe in 15 isolates of which 14 were also RIF resistant by DST. In all 15 of these RIF resistant isolates (by MTBDR*plus*) MLPA identified INH resistance, but not RIF resistance. The concordance of the detection of MDR-TB between all methods is shown in FIGURE 2.

**Figure 2.**
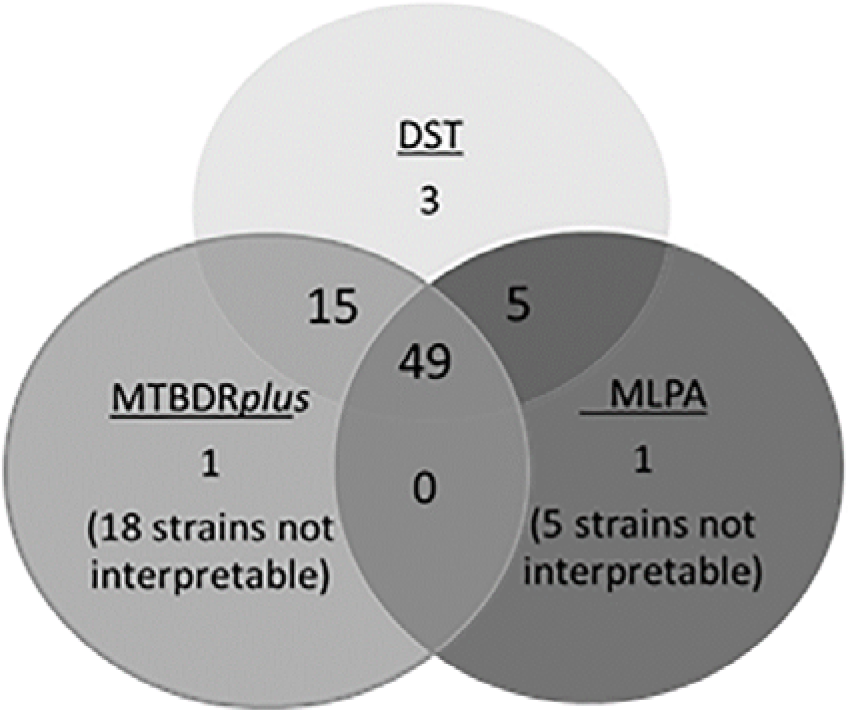
Concordance between all methods used to determine MDR-TB. For the comparison results from all methods obtained for all strains were used. Numbers indicate strains identified by a single method or by multiple methods (overlapping circles).

Eighty-two strains were screened for second line drug resistance by DST, including 74 M(X)DR-TB strains and eight selected on the basis of poor clinical response. Among these 82 strains eight (8/82, 9.6%) were resistant to KAN, and OFX by DST and were thus XDR-TB. Additionally, DST identified capreomycin resistance in four strains, one of which was also resistant to PAS. All 399 isolates were screened for second line drug resistance by MLPA (TABLE A4). MLPA detected OFX resistance in 17/399 isolates screened; the gyrA-A90V mutation in eight strains (six of which were OFX resistant by DST); the gyrA-D94G mutation in nine strains (eight of which were OFX resistant by DST). Sequencing of the *gyrA* gene was performed on one strain identified as XDR-TB by DST and MLPA and confirmed the presence of the gyrA-D94G mutation detected by MLPA. Sequencing showed that the two quinolone resistant strains identified by DST, but not by MLPA, did not carry a mutation in *gyrA.* MLPA detected the rrs-1401 mutation associated with resistance to KAN/AMK/Capreomycin in 10 of the 399 isolates (three of which were XDR, and four MDR by DST). In one of the 399 isolates, strain 12-15893, MLPA detected a mutation in the *eis* gene, this strain was XDR by DST (TABLE A4).

Of the 394 (98.7%) strains with an interpretable MLPA profile, 248 (62.9%) were members of the Euro-American lineage (FIGURE 3). Of these 248 strains, 88 were further sub-classified as LAM (62/394; 15.7%), Haarlem (23/394; 5.8%), CAS (2/394; 0.5%), or X lineage (1/394;0.2%). The second largest group was Beijing, 140/394 (35.5%) strains. MLPA subdivided the Beijing strains into Beijing K1 (95/394; 24.1%), Beijing V+/CHIN+ (43/394;10.9%), Beijing SA-/CHIN-,or Beijing V-(1 and 1 each 0.3%). MLPA profiles of 6/394 (1.5%) samples showed the presence of multiple lineage markers assumed to represent mixed infections.

**Figure 3.**
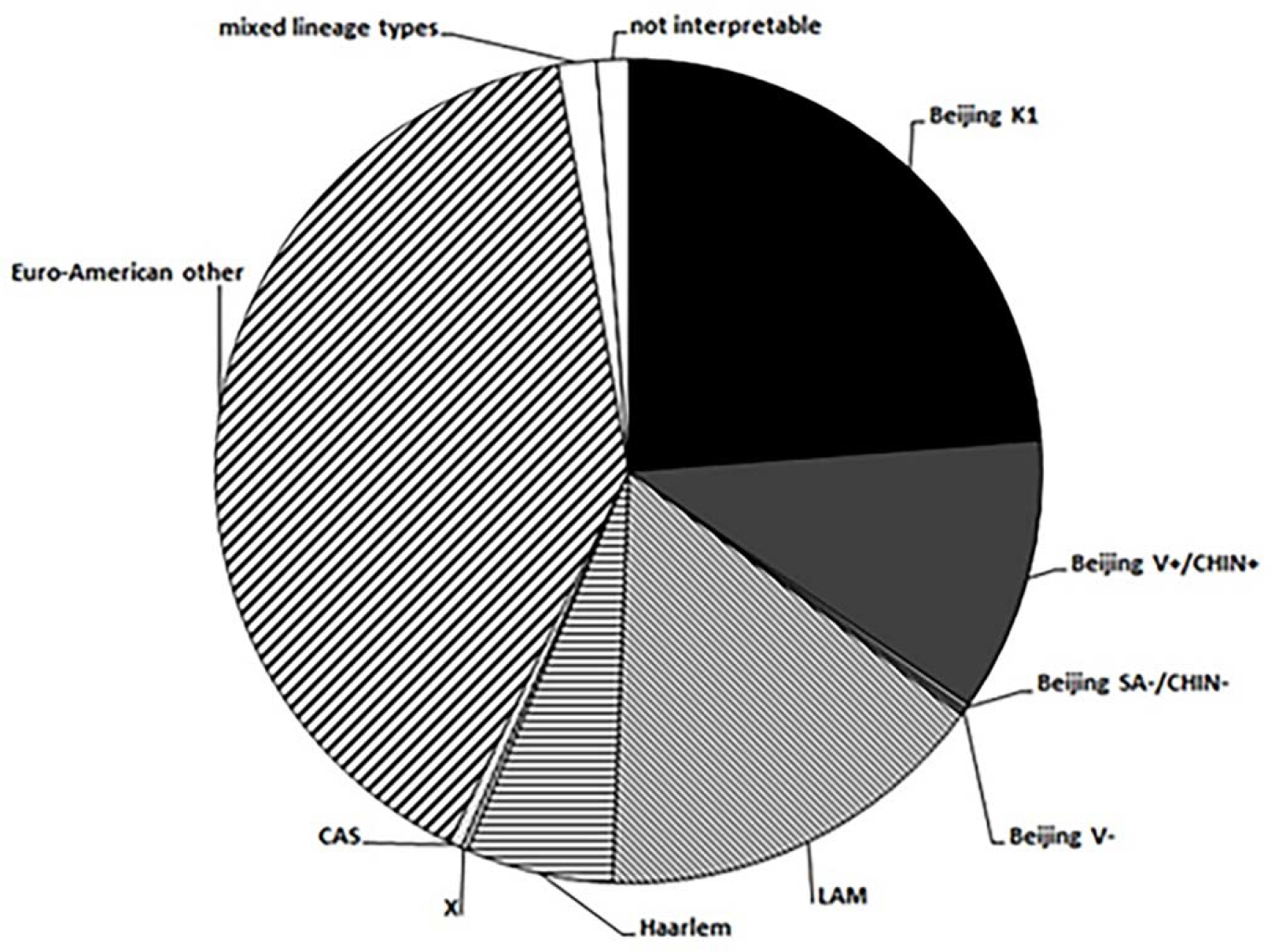
Mycobacterium t 455 uberculosis lineage diversity in 399 cultured clinical isolates from pulmonary TB patients in Tbilisi, Georgia between 2012–2013. Beijing lineage (solid): Beijing K1 lineage (n=95), Beijing V+/CHIN+ (n=43), Beijing SA-/CHIN-(n=1), Beijing V-(n=1); Euro-American lineage (patterned): LAM (n=62), Haarlem (n=23), X lineage (n=1), CAS (n=2), Euro-American other (n=160); Mixed lineage types/notinterpretable (white): mixed lineage types (n=6), not interpretable (n=5).

Combining the data above revealed 148/248 Euro-American strains (60%) were pan-susceptible by DST and 52/248 Euro-American strains (52/248, 21%) were monoresistant to streptomycin (TABLE 2). Only 3.6% of the Euro-American strains (9/248) were MDR-TB (non XDR-TB) of which five were resistant to all tested first line drugs. In contrast 45% (63/140) of all Beijing strains identified were MDR-TB (eight XDR-TB) of which 43% (60/140) were resistant to all tested first line drugs by DST. Of the remaining Beijing strains 54 (54/140, 38%) were pan-susceptible, eight (8/140, 6%) were resistant only to streptomycin, and 15 (15/140, 11%) were resistant to INH and/or S and EMB (TABLE 2).

**TABLE 2.** MLPA lineage profiles for 399 Georgian isolates stratified according to their DST profile.

| Lineage type by MLPA | (% of total) | Drug Susceptibility Testing, SIRE |  |  |  |  |  |  |  |
| --- | --- | --- | --- | --- | --- | --- | --- | --- | --- |
|  |  | RRRR | RRRS | SRRS | RRSS | SRSS | RSSS | RRSR | SSSS |
| Total Beijing (n=140) | 35.1 | 60 (43) | 2 (1) | 1 (1) | 7 (5) | 4 (3) | 8 (6) | 4 | 54 (38) |
| Beijing K1 (n=95) | 23.8 | 27 |  | 1 | 2 | 4 | 7 | 3 | 51 |
| Beijing V+/CHIN+ (n=43) | 10.7 | 32 | 2 |  | 5 |  | 1 | 1 | 2 |
| Beijing SA-/CHIN- (n=1) | 0.0 |  |  |  |  |  |  |  | 1 |
| Beijing V- (n=1) | 0.0 | 1 |  |  |  |  |  |  |  |
| Total Euro-American (n=248) | 62.2 | 5 (2) | 4 (2) |  | 16 (6) | 18 (7) | 52 (21) | 2 (1) | 149 (60) |
| LAM (n=62) | 15.5 | 1 | 1 |  | 5 | 7 | 6 |  | 41 |
| Haarlem (n=23) | 5.7 | 2 |  |  | 1 | 3 | 1 | 1 | 15 |
| X (n=1) | 0.0 |  |  |  |  |  |  |  | 1 |
| CAS (n=2) | 0.0 |  |  |  |  |  |  |  | 2 |
| Euro-American other (n=160) | 40.1 | 2 | 3 |  | 10 | 8 | 45 | 1 | 90 |
| mixed lineage types (n=6) | 1.5 |  | 1 |  |  |  | 1 |  | 4 |
| non-interpretable/ NTM (n=5) | 1.2 | 1 |  |  | 1 |  | 2 | 1 |  |
| Total (N = 399) | 100 | 66 | 7 | 1 | 24 | 22 | 63 | 7 | 207<sup>a</sup> |

SIRE = streptomycin/ isoniazid/ rifampicin/ ethambutol. a, for one strain the DST result for EMB was not reported (SSSX); SRSR: one Euro-American other isolate; SSSR: one LAM isolate. Numbers in brackets indicate percentages of drug resistance within one MTB lineage.

In this unselected set of isolates MDR-TB cases were detected in 36 of 289 (12.2%) new cases and 38 of 100 (38.0%) retreatment cases. These patient characteristics were considered with respect to resistance profile and *Mtb* lineage and correlations were analyzed using univariate and multivariate regression analysis (TABLE 3) and visualized in a Sankey diagram (FIGURE 4).

**Figure 4.**
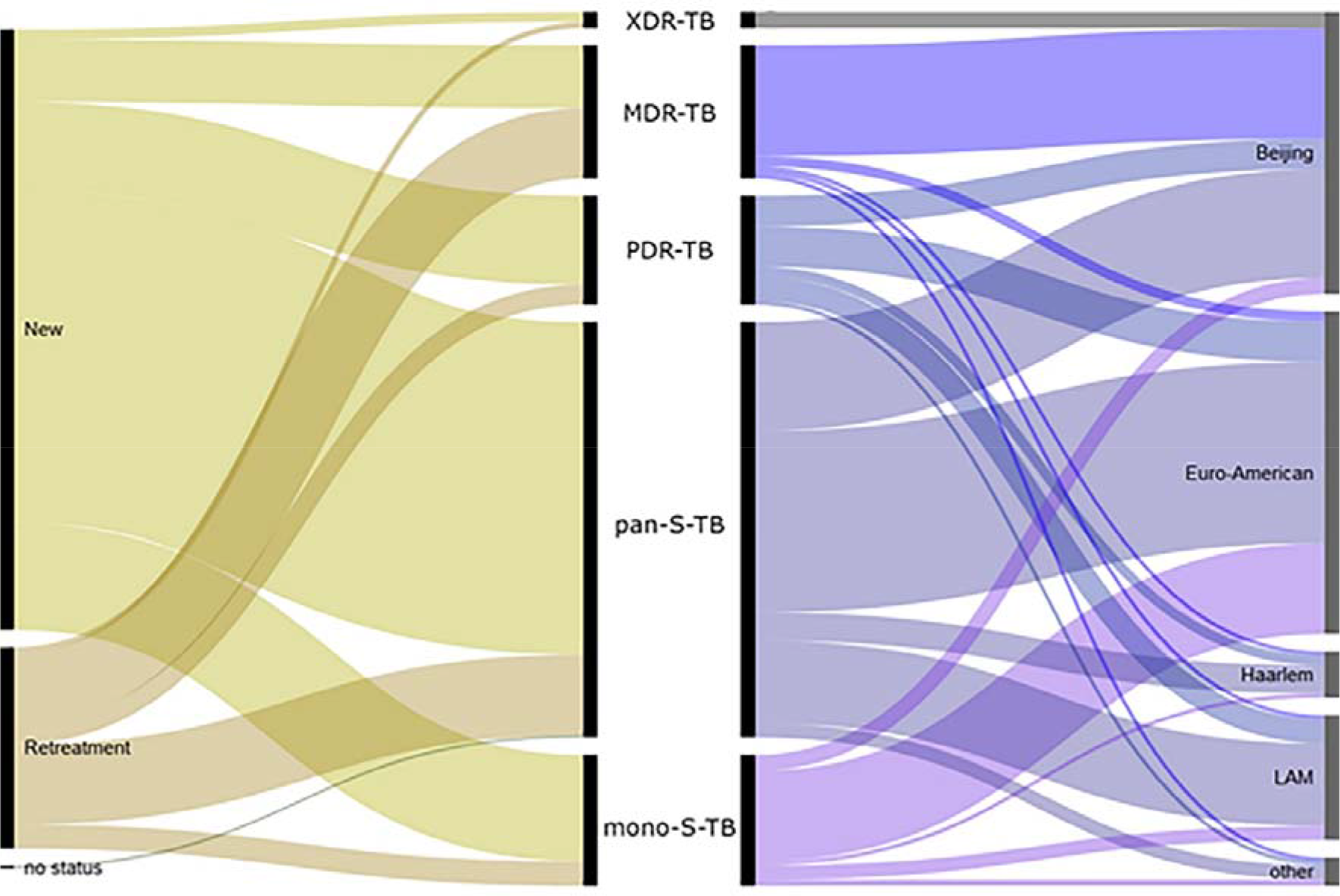
Sankey diagram showing the relationships between treatment history, drug susceptibility results and Mtb lineage of all 399 samples tested by DST and MLPA. XDR-TB are exclusively from the Beijing lineage, MDR-TB are overrepresented in the Beijing lineage (85% Beijing), 38% of retreatment cases were MDR-TB. New (n=298), Retreatment (n=100), no status (n=1); XDR-TB (n=8), MDR-TB (n=66), polydrug-resistant (PDR)-TB (n=54), pan susceptible (pan-S)-TB (n=206), mono-STR (mono-S)-TB (n=63) and other (n=2); Beijing (n=140), Euro-American (n=160), Haarlem (n=23), LAM (n=62), Other (n=6, suspected mixed strains) and (n=5, not interpretable) and (n=2, CAS lineage) and (n=1, X lineage). The Sankey diagram was designed with the webtool RAW (http://app.raw.densitydesign.org/).

**TABLE 3.** Estimated effect of patient and strain characteristics on the odds of a TB patient having MDR-TB (logistic regression)

| Variables | n | univariate analysis |  | multivariate analysis (n=387) |  |
| --- | --- | --- | --- | --- | --- |
|  |  | OR (95% CI) | p-value<sup>a</sup> | OR (95% CI) | p-value |
| Age | 393 | 0.98 (0.97 to 1.01) | 0.222 |  |  |
| Male | 392 | 1.02 (0.57 to 1.87) | 0.940 |  |  |
| Strain (vs. Euro-American<sup>b</sup>) | 387 |  | <0.001 |  | <0.001 |
| Beijing |  | 21.63 (10.30 to 45.54) | <0.001 | 20.12 (9.41 to 43.04) | <0.001 |
| Treatment history (vs. new) | 392 |  |  |  |  |
| Retreatment |  | 4.59 (2.68 to 7.86) | <0.001 | 3.95 (2.08 to 7.54) | <0.001 |

(a) Wald test of association for individual odds ratio (OR), log likelihood ratio test for overall test of significance (categorical variables); (b) including Haarlem/LAM/CAS/X lineage and Euro-American other.

## DISCUSSION

Here we evaluated the feasibility, performance and potential information obtainable by introducing and performing a SNP-based molecular assay for genotyping *Mycobacterium tuberculosis* at the NRL of the NCTLD in Tbilisi, Georgia. SNP based characterization was possible for all but five of 399 isolates. Linking this data with the routine DST and patient information allowed an initial assessment of the dynamics of the TB epidemic in Georgia. There were striking differences between the risk of an MDR phenotype and specific *Mtb* lineages.

This study has limitations. Our samples size represents only approximately 10% of all notified TB cases for the year 2012 (23). The MLPA assay and the standard methods were not performed on the same sample. In 68.2% (272/399) of all samples tested the MTBDR*plus* assay was performed directly on sputum whereas the MLPA assay was exclusively performed on cultured isolates. The MLPA assay was performed on site by the local laboratory staff for monitoring purposes at the end of the month and not as a routine tool such as the MTBDR*plus* assay which is performed on a daily basis. Minor problems were experienced, mainly related to the stability/functionality of the Luminex MAGPIX device but none of these prevented the assay from being performed always yielding good quality data. However the analysis and interpretation of the data required remote support. Either straight forward data analysis and interpretation or timely online support is a prerequisite for any molecular tool to be used in a routine diagnostic lab. Optimizing the use of data generated for real time monitoring rather than remote analysis is desirable.

The MLPA assay targets only the most common resistance associated mutations. For this reason it did not detect a proportion of RIF resistant strains detected by the reference standards. Accordingly, calling of an MDR-TB genotype by the MLPA alone lacked sensitivity. The currently MLPA cannot replace DST combined with line probe assays for clinical management, but sequence-based drug-resistance testing could conceivably achieve this (32, 33). However, a high specificity was obtained for the detection of M(X)DR-TB by MLPA. Of the eight XDR-TB strains identified by DST, resistance to AMK/KAN/capreomycin was identified in only half of the samples by MLPA. Mutations outside of the hot spot region of the *rrs* gene may account for the numbers of resistant phenotypes. Mutations in the *eis* gene have been associated with resistance to KAN (34–36). In this study MLPA identified a single isolate with an XDR phenotype that also carried a mutation in the *eis* gene and was a Beijing K1 strain.

Some of the discrepancies observed between the three methods of screening for drug resistance (FIGURE 2) may have been due to the presence of multiple resistance genotypes (37) a fact supported by the observation that a significant minority (9.8%) of the MLPA resistance calls were intermediate. The current study thus provides additional evidence supporting the interpretation of intermediate MLPA values for resistance associated mutations as described previously (24, 28). In this study, 43 intermediate values were obtained that could be compared to a reference standard. For 31(72.1%) of these intermediate values resistance was detected by the reference standard. Thus intermediate MLPA values are highly suggestive of heteroresistance. Mixed resistance genotypes are often observed in high MDR settings (37). The relative contribution to mixed genotypes as a result of cross infection with resistant genotypes or resistance amplification deserves further study.

Association of resistance and patient characteristics to the genotypes: Of all M(X)DR-TB detected by DST 85% (63/74) were strains of the Beijing lineage. The MLPA is able to sub-delineate Beijing into five sub-lineages (26). Two Beijing sub-lineages (Beijing V+/CHIN+ and Beijing K1) accounted for 84% of all the MDR-TB identified. Additionally 29% (28/95) of all Beijing K1 lineage strains and 79% (34/43) of all Beijing V+/CHIN+ strains were MDR. All XDR-TB isolates identified were members of the Beijing lineage (Figure 4). MIRU-VNTR typing (TABLE A4) revealed that 18 of the 34 MDR-TB Beijing V+/CHIN+ strains belonged to the MLVA 15–9 type 100–32 and all 28 MDR-TB Beijing K1 strains belonged to the MLVA 15–9 type 94–32. Both 100–32 and 94–32 represent epidemic MDR-TB cluster types (11, 38) which have been previously identified in Georgia (24). The 100–32 cluster was formed exclusively by Beijing V+/CHIN+ lineage M(X)DR-TB strains, whereas the 94–32 cluster was formed by strains of the Beijing K1 and Beijing V+/CHIN+ lineage with various drug resistance profiles except streptomycin mono-resistance.

Although an MDR phenotype was associated with retreatment, *Mtb* lineage was much more strongly associated in this data set. After univariate analyses individuals infected with a Beijing strain had 20-fold higher odds (21.63, 95% CI 10.30 to 54.54) of being MDR-TB than individuals infected with a Euro-American strain; whereas retreatment patients had a 4-fold higher odds of being infected with an MDR-TB (4.59; 95% CI 2.68 to 7.68) (TABLE 3 and FIGURE 4). Multivariate analysis confirmed that the effects of Beijing strain and retreatment were independent (TABLE 3).

High *Mtb* cluster rates among previously hospitalized HIV patients co-infected with XDR-TB(10) and reported TB infection among hospital workers suggests nosocomial transmission as a main factor facilitating transmission of drug resistant strains. A high incidence of MDR-TB strains in penitentiary systems (39), transmission of these strains in the community through released inmates, prison staff and visitors (40) might also facilitate spread of MDR-TB strains in high burden MDR-TB countries.

Of all strains with any drug resistance identified by DST 32.0% (63/192) were streptomycin monoresistant. The Euro-American lineage was over represented in the streptomycin-monoresistant strains, 85.2% 52 out of 61 were from the Euro-American lineage. Of these 52 streptomycin monoresistant isolates 13 (25%) belonged to the MLVA 15-9 type 769-15. This MIRU type was identified in Georgia and named Georgia H37Rv-like (5, 24); indicating that a proportion of the ancestors of the circulating Euro-American strains ‘”witnessed” streptomycin and their progeny are still circulating.

Rapid molecular testing has been recently shown to significantly decrease the time to initiation of appropriate MDR-TB treatment in Georgia (3, 4). Synthesis of the bacterial lineage data with available DST and patient characteristics here strikingly demonstrated that multidrug resistance is significantly more associated with the Beijing lineage thana previous history of TB treatment in Georgia. To objectively measure the relative contribution of cross infection versus resistance amplification in diverse settings we suggest that the ratio of risk of MDR-TB associated with retreatment versus bacterial lineage is an interesting metric which could be used to express the contribution of resistance generation vs transmission, and should be further explored.

Combining resistance and genotyping data with patient characteristics will become increasingly practical to implement. A combined approach of spatial and molecular with classical epidemiology to study the transmission of (MDR)-TB has been shown to be feasible in Georgia. Infection control as well as treatment and patient management could benefit from additional knowledge of the infecting *Mtb* lineages (41, 42) and aid the identification of outbreak strains that might otherwise be missed (43). Most strikingly in this pilot implementation, when the genotyping patient data and susceptibility data were combined it was observed that a patient infected with a Beijing strain had 20-fold higher odds of being MDR-TB than a patient infected with a Euro-American strain. Interestingly a retreatment case of TB had “only” a 4-fold higher odds of being MDR-TB than a primary case. Monitoring these associations could help to understand the local transmission dynamics and identify areas where resources should be targeted. TB control programs can directly use genotyping data, and in the future WGS data, to rationally develop, adapt and prioritize infection control efforts but only if it is rapidly integrated with patient and bacteriological data: Such a goal is becoming increasingly necessary but also realistic.

## Acknowledgements

The TB-MLPA as described in the text is commercially available as the TB-SNPID assay, distributed via Beamedex, Orsay, France www.beamedex.com. KIT BR has a financial interest in the assay.

## Supplemental files

TABLE A1. Performance parameters of the MLPA detecting molecular resistance to first and second line drugs compared to conventional DST as the reference standard.

TABLE A2. Performance parameters of the MLPA detecting molecular resistance to INH and RIF compared to GenotypeMTBDRplus as the reference standard.

TABLE A3. Correlation between drug resistance identified by MLPA and MTBDRplus

TABLE A4. Results obtained for drug resistance for all isolates by sputum microscopy, DST for first line and second line drugs, GenotypeMTBDRplus, MLPA and MIRU-VNTR.

